# Blood meal analysis of tsetse flies (*Glossina pallidipes*: Glossinidae) reveals higher host fidelity on wild compared with domestic hosts

**DOI:** 10.1101/692053

**Authors:** Manun Channumsin, Marc Ciosi, Dan Masiga, Harriet Auty, C. Michael Turner, Elizabeth Kilbride, Barbara K. Mable

**Affiliations:** Faculty of Veterinary Medicine, Rajamangala University of Technology Tawan-Ok, Chonburi, 20110, Thailand; Institute of Molecular, Cell and Systems Biology (MCSB), University of Glasgow, University Place, Glasgow G12 8QQ, UK; International Centre of Insect Physiology and Ecology (ICIPE), P.O. Box 30772, 00100 Nairobi, Kenya; Institute of Biodiversity, Animal Health and Comparative Medicine (BAHCM), Graham Kerr Building, University of Glasgow, University Place, Glasgow G12 8QQ, UK; Institute of Infection Immunity and Inflammation (III), University of Glasgow, University Place, Glasgow G12 8QQ, UK

**Keywords:** blood meal, tsetse flies, Trypanosomes, parasites, host, host population structure, African trypanosomiasis

## Abstract

Changes in climate and land use can alter risk of transmission of parasites between domestic hosts and wildlife, particularly when mediated by vectors that can travel between populations. Here we focused on tsetse flies (genus *Glossina*), the cyclical vectors for both Human African Trypanosomiasis (HAT) and Animal African Trypanosomiasis (AAT). The aims of this study were to investigate: 1) the diversity of vertebrate hosts that flies fed on; 2) whether host feeding patterns varied in relation to type of hosts, tsetse feeding behaviour, site or tsetse age and sex; and 3) if there was a relationship between trypanosome detection and host feeding behaviours or host types. Sources of blood meals of *Glossina pallidipes* were identified by sequencing of the mitochondrial cytochrome b gene and analyzed in relationship with previously determined trypanosome detection in the same flies. In an area dominated by wildlife but with seasonal presence of livestock (Nguruman), 98% of tsetse fed on single wild host species, whereas in an area including a mixture of resident domesticated animals, humans and wildlife (Shimba Hills), 52% of flies fed on more than one host species. Multiple Correspondence Analysis revealed strong correlations between feeding pattern, host type and site but these were resolved along a different dimension than trypanosome status, sex and age of the flies. Our results suggest that individual *G. pallidipes* in interface areas may show higher feeding success on wild hosts when available but often feed on both wild and domesticated hosts. This illustrates the importance of *G. pallidipes* as a vector connecting the sylvatic and domestic cycles of African trypanosomes.

## Introduction

In sub-Saharan Africa, changes in land use increase encroachment of domestic livestock into areas that are primarily managed to conserve wildlife. This increases risks that livestock will be exposed to a wider range of parasites, with potentially important consequences for disease burden and control. Wildlife can represent ‘reservoir communities^31,71^ for multi-host pathogens that could spill-over into domesticated animals. Domesticated animals infected by wildlife pathogen could in turn show more severe disease, given limited opportunity for host-pathogen coevolution in novel hosts. This could be particularly true for vector-mediated transmission, where movement of the vectors could facilitate parasite sharing across interface areas, even if fences are used to reduce contact between domestic and wild hosts.

One particularly complex system where this could be important to understand is trypanosome-mediated diseases transmitted by tsetse flies in Africa. Although there are multiple species of tsetse flies that can transmit multiple species of trypanosomes, *Glossina pallidipes* is the most economically important species in East Africa ^21^, because it is the main vector of Animal African Tryanosomiasis (AAT) and it is also a vector of Human African Trypanosomiasis(HAT). Wild animals have been reported as reservoir hosts both for AAT^3,57^ and HAT^25,26,34,73,74^ but the extent of transmission across the wildlife-livestock interface remains unclear.

Tsetse flies (genus *Glossina*) are generalist blood-feeders on a wide variety of vertebrate host species, including mammals, reptiles and birds^72^. Importantly, both male and female tsetse feed throughout their lifetimes. There is thus high potential for vector-mediated connection between parasite sylvatic and domestic cycles in wildlife-livestock interface areas if tsetse flies take meals from different host species at each feeding opportunity. However, the likelihood that an individual tsetse feeds on different types of hosts where they occur sympatrically, compared to feeding predominantly on a single species, has not been clearly established and so the relative risks of increased trypanosome infections in livestock living near wildlife remains a critical gap in knowledge^6^. Although three trypanosome species are traditionally associated with the disease in livestock (*T. brucei*, *T. congolense*, and *T. vivax*), a higher diversity has been identified in wildlife^4^, which could potentially increase risks of disease if transmission from wildlife to domesticated animals is common.

Few studies have attempted to combine investigation of host-feeding patterns of individual flies, trypanosome infection, and intrinsic factors of tsetse flies distributed in different regions. Identification of hosts through blood meal analyses is a highly useful tool that has been used to predict host preferences and feeding behaviours across a wide range of vectors^24,28,39^ A commonly used approach has been to use polymerase chain reaction (PCR)-based techniques to amplify and sequence host DNA from blood meal contents in the guts of fed flies. This has largely been based on mitochondrial genes due to their high copy number and the extensive databases available due to their use as universal markers for DNA barcoding^33,42,60^. For example, in the Serengeti ecosystem in Tanzania, which holds a high number and wide range of wild animals, an investigation of blood meal composition in tsetse flies based on sequencing of the cytB gene compared to surveys of host density revealed strong preferences for particular wild hosts, which varied by species of fly (G*. swynnertoni* vs *G. pallidipes*^5^. This clearly demonstrated the value of relating feeding patterns to the diversity of hosts present. However, trypanosome prevalence was not quantified in these studies and domestic hosts were not present in the study area; so, relative host preferences for wildlife compared to livestock was not determined.

Feeding activity, where individual flies feed consecutively on different types of hosts, could alter relative risk of transmission of trypanosomes. More frequent feeding might occur, for example, if flies are disrupted while feeding or if they abandon a host that they perceive to be unsuitable or that shows defensive behaviour^63,68^. The dominance of nonpreferred hosts in a particular geographic area could thus result in more frequent host switching and so increased rates of multiple feeding and potentially higher exposure to a diverse range of parasites. In East Africa, *G. pallidipes* is widespread and has been demonstrated to feed on a wide range of hosts, including bovines^19,20,54,58,69^, suids^11,58^, elephants^54^, antelopes^1^ and cattle^54^. Warthogs, bushbuck and African buffalo have been suggested as the preferred hosts^13,19,43,45,54,58^ but this varies by geographic region^5,22,54,65,66^ and relative preference for domestic and wild hosts has not been specifically assessed.

In a previous study, we established that the prevalence of trypanosomes among tsetse flies in two regions of Kenya (Nguruman and the Shimba Hills) showed complex relationships with geographic location, tsetse specific factors (age, sex and fly species), species of trypanosome and the presence of an endosymbiont^17^. The main aim of the current study was to assess whether some of the variation in the detection of trypanosomes across sites could be explained by differences in host feeding patterns. Specifically, we aimed to determine: 1) the diversity of vertebrates tsetse fed on at sites where different types of host were present; 2) whether host feeding patterns varied in relation to type of hosts or intrinsic tsetse factors (i.e. age and sex); and 3) if there was a relationship between trypanosome detection and host feeding patterns, host types or tsetse-specific factors.

## Methods

### Sampling and tsetse fly characterisation

The *G. pallidipes* samples are described in Channumsin *et al*.^17^, where details of the sampling strategy are provided (see Extended data 1^18^ for locations of the traps). NG2G traps baited with acetone and cow urine^15^ were used for collecting tsetse flies from three sites that differ in anticipated levels of relative abundance of livestock and wildlife, with the sampling effort (number of traps) determined by the relative abundance of flies in the area. Two sites were sampled in the Shimba Hills National Reserve (Kwale County, in the coastal region of Kenya), which is a relatively small (250 km^2^) protected area separated from surrounding agricultural areas by a wildlife fence. There is extensive habitat for tsetse flies, including on the park boundaries. Buffalo Ridge is within the fenced wildlife protected area in the middle of a thicket forest, where many tourists visit all year, while Zungu Luka has a woodland type of vegetation, and is located on the border of the park close to a permanently human-inhabited rural area with resident livestock. In contrast, the Nguruman region contains lowland woodland patches surrounded by open savannah; habitats, which have been found to host a large number of *G. pallidipes* and *G. longipennis*^16^. The sampling site (Mukinyo) is at the border of the Olkiramatian group ranch, which is a wildlife conservancy without fences, where the distribution of domestic and wild tsetse hosts overlap when livestock are grazed in the area but there is no permanent human settlement close by.

Characteristics of the flies and presence of trypanosomes were previously determined by Channumsin *et al*.^17^. Sex and species of flies were determined based on morphological characters. Age was estimated based on a wing fray score where increased damage indicates increasing age^37^., Whole flies were preserved in 95% ethanol and stored at −20°C. Presence of trypanosomes in mouth parts and proboscis of the flies collected was determined using general primers targeting the ITS-1 region of the rDNA array (CF: 5’ CCGGAAGTTACCGATATTG 3’ and BR: 5’ TTGCTGCGTTCTTCAACGAA 3’^56^, that allow identification of trypanosome species based on size of amplicons, as described in Channumsin *et al.*^17^. Although multiple species of tsetse were used in the previous study, here we focused on individuals identified morphologically as *G. pallidipes* (N = 577) and screened trypanosomes in DNA that had been extracted from abdomens. All flies sampled were used, rather than selecting individuals that had appeared to have fed recently.

### Identification of diversity of hosts and feeding patterns from *G. pallidipes* blood meals

We used primers developed by Kocher *et al*.^40^ targeting a 359 bp fragment of the mitochondrial gene cytochrome B (cytb) gene in mammals (Cb1: 5’ CCATCCAACATCTCAGCATG ATGAAA 3’ and Cb2: 5’ GCCCCTCAGAATGATATTTGTCCTCA 3’) which enabled direct comparison with two previous studies^4,54^ and because they showed more reliable amplification in a pilot study^75^ than primers targeting the mitochondrial cytochrome C oxygenase 1 (CO1) gene (VF1d-t1 and VR1d-t1)^36^. During processing for DNA extractions, in order to reduce risk of contamination, the dissected tissues were cleaned 2-3 times with 95% ethanol, then left to dry, before moving to new individual microtubes with liquid nitrogen for sample crushing and DNA extraction using DNeasy® blood and tissue kits (Qiagen Inc., Paisley, UK). PCR cycling was carried out in 25 μl reaction mixtures containing: 1X PCR buffer; 0.2 mM dNTP mixture; 1.5 mM MgCl_2_ (Thermo Scientific); 0.5 μM of each primer; 1 unit of *Taq* DNA polymerase (Invitrogen Inc, Carlsbad, CA., USA); and 2 μl tsetse abdomen DNA template. Samples were pre-heated at 94°C for 5 min, denatured at 94°C for 30 sec, annealed at 55°C for 45 sec, then extended at 72°C for 30 sec, with 35 cycles of the amplification and a final extension at 72°C for 10 min^54^. PCR products were visualised using 1.5% UltraPure™ Agarose gels (Invitrogen, Paisley) with 2% Ethidium Bromide (Invitrogen, Paisley) in 1X TBE buffer (108 g of Tris Base, 55 g of Boric acid and 40 ml of 0.5 M EDTA). Results were visualised and analysed on a gel documentation system (UVIpro Platinum, UVITEC, Cambridge, UK or GeneDoc, BioRad Inc, UK).

PCR products of the expected size (359 bp) yielding ≥ 20 ng were cleaned using ExoSAP-IT PCR Clean-up Kits (GE Healthcare). In cases where the yield of PCR products was lower than this threshold, multiple PCR products were concentrated and QIAquick Gel Extraction Kits (Qiagen Inc, Paisley, UK) were applied to extract the PCR products from agarose gels. All purified samples were sent for Sanger sequencing in both forward and reverse directions, using the Sequencing Service at the University of Dundee (MRC I PPU, School of Life Sciences, University of Dundee, Scotland, https://www.dnaseq.co.uk) using Applied Biosystems Big-Dye Ver 3.1 chemistry on an Applied Biosystems model 3730 automated capillary DNA sequencer.

Base-calling was manually corrected, sequences were aligned and consensus sequences for forward and reverse primers for each individual generated using Sequencher version 5.3 (Gene Codes Corporation, Ann Arbor, MI USA). The Basic Local Alignment Tool (BLASTn)^2^, was used to identify the closest matching sequences in the GenBank database to determine the host identity of each consensus sequence. Chromatographs with only single peaks based on direct sequences were classified as “single host feeding”. Sequences that were still clearly readable but showed more than one peak at multiple positions were classified as “multiple host feeding”. While the difficulty of resolving the phase of genetic variants precluded identification of all hosts from direct sequencing of multiple-peak products, the dominant host was determined based on BLASTn analysis of the most prominent peaks.

Host-feeding patterns were confirmed in a subset of samples by cloning using TOPO®-TA Cloning Kits (Invitrogen, UK), with at least six plasmids of each sample sent for sequencing, after purifying using QIAprep Spin Miniprep Kits (Qiagen Inc, Paisley, UK). Ten samples whose chromatographs showed double or triple peaks at single positions in the direct sequences were cloned to confirm that multiple peaks were due to feeding on multiple host species rather than poor quality sequences (five flies from Buffalo Ridge; three from Zungu Luka; two from Mukinyo). An additional seven samples that appeared to have fed on single hosts but with some ambiguous peaks were also cloned and sequenced (five from Buffalo Ridge and two from Zungu Luka).

To enable of assessment of variation in the type of hosts fed on across sites, dominant hosts resolved were classified as “domestic” (including livestock or companion animals), “human”, or “wild”.

In order to assess infraspecific diversity in hosts fed on across the sites, sequences were first exported to Se-Al version 2.0^61^ to manually align and prune sequences to the same length. DNAsp version 5.0^47^ was then used to resolve variants into unique haplotypes within host species. Minimum spanning networks were plotted using PopArt^46^, to indicate relative frequencies of host haplotypes across the sampling sites.

### Variables influencing tsetse feeding behaviour

Generalised linear models (using the glm function, as implemented in the lme4 package^8^ using R version 4.0.2^67^ were used to test whether variation in feeding behaviour of the flies (single vs multiple hosts, modelled as a binary response variable) was influenced by the type of the dominant host (domestic, human, or wildlife), tsetse sex and age (sex as a continuous variable based on awing fray score averaged across the two wings of an individual), or site (Buffalo Ridge, Zungu Luka, Mukinyo). Interactions between the type of host with sex, age and site were also considered in the full model. Model selection was performed using likelihood ratio tests to find the minimum model that best explained the data. Odds ratios were calculated from the coefficients of the final model using the “oddsratio” package in R^64^. To check the appropriateness of the binomial model, overdispersion was assessed by checking that the ratio of the residual deviance to the degrees of freedom in the final model was below 1. The fit of the final model was assessed by McFadden’s pseudo-R^2^, defined as 1 – LL(final model)/LL(null model), where LL = log likelihood^52^.

### Prevalence of *Trypanosoma spp.* in relation to *G. pallidipes* feeding patterns

A similar statistical approach was used to test whether the presence of trypanosomes was explained by host type or feeding behaviour while considering possible influences of fly sex and age, or site based on conclusions from our previous study^17^. Since we were specifically interested in whether tsetse feeding behaviour affected trypanosome detection, pairwise interactions were considered between the type of host and the feeding pattern with age, sex and sampling site of the flies. Model selection and fit were performed as described for the feeding pattern models. Given the wide range of hosts that the flies feed on, the influence of particular host species on trypanosome prevalence was considered only qualitatively.

To specifically visualise whether feeding patterns or dominant host types were related to trypanosome prevalence when accounting for geographic location and tsetse sex and age, Multiple Correspondence Analysis (MCA), as implemented in the FactoMineR package (version 1.30^44^) was used. For this analysis, age was considered as a categorical variable by classifying individuals into the following age categories: “young” (wing fray score 1–2.5); “juvenile” (3.0-4.0) and “old” (4.5–6.0) based on the average score between the two wings for each individual. Other variables were: site, presence or absence of *Trypanosoma spp*, sex, feeding pattern (single vs multiple) and dominant host type (domestic, human or wild). Variation along pairs of principal component axes was visualised using “ggplot2()”^27^ in R^67^.

## Results

### Diversity of hosts identified from *G. pallidipes* blood meals

From 573 *G. pallidipes*, 128 flies showed no evidence of a recent blood meal based on lack of amplification products following screening with the Cb1 and Cb2 primers. These samples were excluded from analyses (Table 1). The remaining 445 flies showed amplified products of the expected size, which were sequenced and used to classify feeding status (Table 1; Extended data 2^18^). For 197 of the 247 samples for which dominant hosts could be resolved to the species level, a single amplification product was apparent in the chromatographs; these were classified as having recently fed on a single host. Cloning of seven of these samples confirmed amplification of DNA from only a single host species (Extended data 2 and 3^18^). The chromatographs for 53 samples clearly showed multiple peaks that could be confidently attributed to feeding on multiple hosts rather than poor sequence quality and the dominant host could be resolved through BLASTn analysis of the strongest peaks. This represented 37% (Buffalo Ridge), 31% (Zungu Luka) and 51% (Mukinyo) of the samples screened at the three sites (Table 1). Cloning of 10 of these PCR products confirmed amplification of DNA from more than one host, with up to four different host species identified in single flies (Extended data 3^18^).

**Table 1.** Summary of blood meal analysis results based on direct sequencing. Cytb negative samples were classified as “unfed flies” but were not considered in the analyses since they could represent lack of amplification rather than lack of feeding. Single host feeding refers to cases where the cytb sequence had only single chromatograph peaks. Multiple host feeding were samples for which cytb was amplified but the sequences showed multiple peaks and the dominant sequence could be identified to species, classified as domestic animals, humans or wildlife. Flies showing strongly amplified cytb PCR products but for which the number or type of host species could not be confirmed due to poor sequencing quality are labelled as “not identified”. The number of flies that tested positive for the presence of trypanosomes is indicated in parentheses. The human samples that were potential contaminants (haplotype 1; Figure 3) were excluded.

| Site | Single Feeding |  |  | Multiple Feeding |  |  | Not identified | “Unfed” | Total Screened |
| --- | --- | --- | --- | --- | --- | --- | --- | --- | --- |
|  | Domestic | Human | Wild | Domestic | Human | Wild |  |  |  |
| Buffalo Ridge | 0 | 1 (1) | 31 (8) | 9 (5) | 9 (3) | 5 (3) | 62 (23) | 33 (10) | 150 (53) |
| Zungu Luka | 1 (1) | 6 (2) | 8 (5) | 12 (7) | 11 (8) | 1 (1) | 60 (31) | 29 (22) | 128 (77) |
| Mukinyo | 1 (1) | 0 | 146 (60) | 0 | 0 | 3 (2) | 76 (22) | 66 (20) | 292 (105) |

We took a conservative approach to classifying feeding patterns: chromatograms of the remaining 198 samples from which amplification products were obtained were not considered of sufficient quality to reliably determine the source of the blood meals; these were classified as “unidentified”. While many of these would likely represent multiple feeding, we wanted to avoid confounding with poor sequence quality so they were classified as fed but not identified (Table 1; Extended data 2^18^); only samples with confident dominant host calls were included in the statistical analyses.

Host composition of blood meals varied across sites (Table 2), with buffalo dominating in the two wildlife protected areas (Buffalo Ridge and Mukinyo) and humans predominating in the site bordering the SHNR (Zungu Luka), where no buffalo feeds were detected (Figure 2). Mukinyo had a wider range of wild hosts identified in blood meals than Buffalo Ridge but elephants, antelope and warthog were found at both sites. Flies from Buffalo Ridge also shared most of the same domestic host species as Zungu Luka, suggesting that flies moved across the fenced interface to feed. Across all sites, only a single fly (from Mukinyo) was confirmed to have fed on domestic cattle.

**Figure 1.**
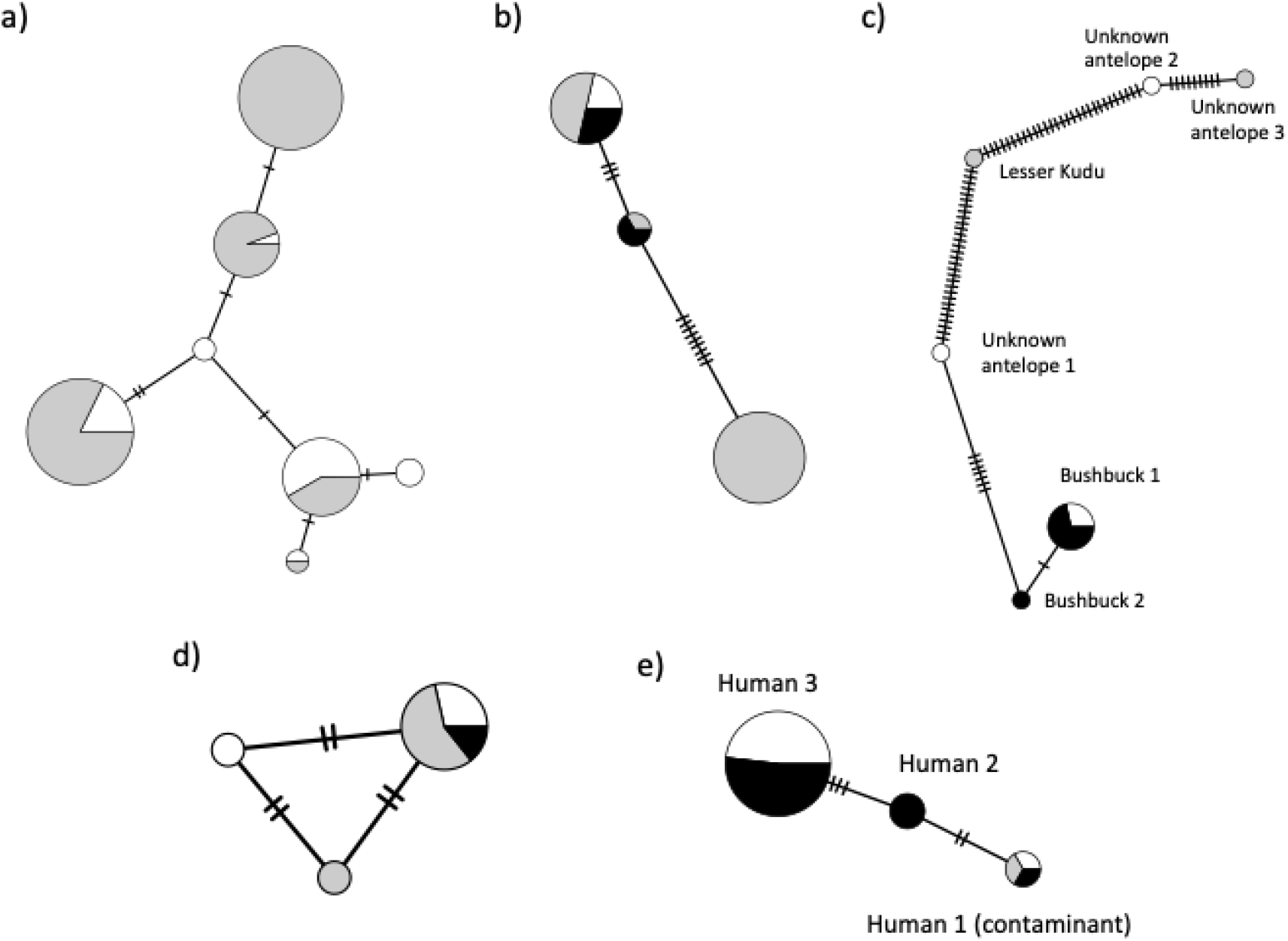
Minimum spanning networks indicating intraspecific diversity and relative frequency of haplotypes between populations from this study for: a) buffalo; b) elephants; c) antelope; d) warthogs; and e) humans. Note that human type 1 matched the ethnic origin of the main investigator (Asian); samples with this haplotype were considered as contaminants and excluded from analyses. The two antelope sequences labelled “unknown” were found only in single clones, with a more dominant host predominating, and had no close match using BLASTn. Circle sizes are proportional to the frequency of each haplotype (see Extended data 4^18^ for values); notches on branches indicate the number of nucleotide substitutions separating haplotypes; colours represent the population of origin (white = Buffalo Ridge; grey = Mukinyo; black = Zungu Luka). Three haplotypes were found in giraffes but they differed by only single nucleotide and were each found in only one or two individuals so they are not shown here.

**Figure 2.**
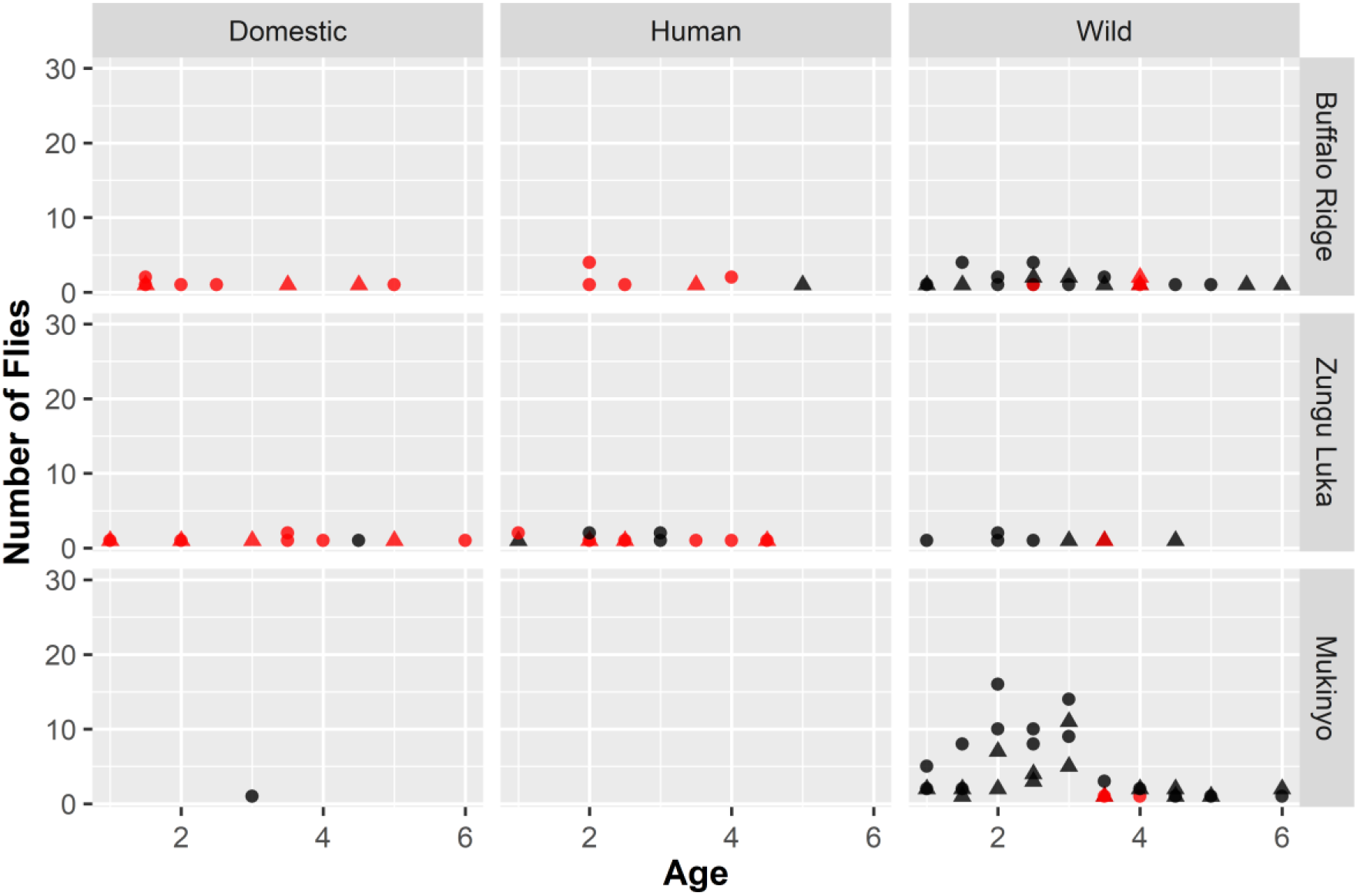
Feeding patterns of flies across sites in relation to their age, sex, and dominant host type (domestic, human, wild). Age of flies was estimated based on average wing fray scores across the two wings or an individual fly, with increasing damage indicating relatively older flies. Feeding patterns were based on whether sequence chromatograms indicated amplification of a blood meal from a single host (black) or more than one host (red); the sex of the flies was determined visually (female = circles; male = triangles). Only values greater than 0 have been plotted. Flies from all age classes at Mukinyo fed predominantly on single wild host species, with no evidence of feeding on humans and only a single mid-age female feeding on a domestic cow. In contrast, feeding on a mixture of domestic and wild hosts was found for all age classes at the Shimba Hills sites, Buffalo Ridge and Zungu Luka and multiple feeding was more frequent than single feeding, except for wild hosts.

**Table 2.** Dominant host species resolved from blood meal analysis *G. pallidipes* sampled from the Shimba Hills (Buffalo Ridge and Zungu Luka) and the Nguruman region of Kenya, based on direct sequencing of cytb. Homozygous amplicons were classified as “single” feeding whereas sequences with multiple peaks were classified as having fed on “multiple” hosts. The dominant host was identified based on BLASTn. The relative abundance of the various host species is expressed as the total % of sequences for which the dominant host could be identified within that site. Species are ordered by relative abundance of wild and domestic hosts.

| Site | Buffalo Ridge |  |  | Zungu Luka |  |  | Mukinyo |  |  |
| --- | --- | --- | --- | --- | --- | --- | --- | --- | --- |
| Host | Single | Multiple | Total % | Single | Multiple | Total % | Single | Multiple | Total % |
| Buffalo | 25 | 3 | 50.9 | 0 | 0 | 0.0 | 105 | 2 | 71.3 |
| Elephant | 2 | 1 | 5.5 | 1 | 1 | 5.1 | 29 | 1 | 20.0 |
| Antelope | 2 | 1 | 5.5 | 6 | 0 | 15.4 | 1 | 0 | 0.7 |
| Warthog | 2 | 0 | 3.6 | 1 | 0 | 2.6 | 5 | 0 | 3.3 |
| Giraffe | 0 | 0 | 0.0 | 0 | 0 | 0.0 | 4 | 0 | 2.7 |
| Hyaena | 0 | 0 | 0.0 | 0 | 0 | 0.0 | 2 | 0 | 1.3 |
| Human<sup>a</sup> | 1 | 9 | 18.2 | 6 | 11 | 43.6 | 0 | 0 | 0.0 |
| Goat | 0 | 7 | 12.7 | 1 | 7 | 20.5 | 0 | 0 | 0.0 |
| Mouse | 0 | 2 | 3.6 | 0 | 4 | 10.3 | 0 | 0 | 0.0 |
| Chicken | 0 | 0 | 0.0 | 0 | 1 | 2.6 | 0 | 0 | 0.0 |
| Cattle | 0 | 0 | 0.0 | 0 | 0 | 0.0 | 1 | 0 | 0.7 |
| Total | 32 | 23 |  | 14 | 25 |  | 147 | 3 |  |

<sup>a</sup> Excluding potential contaminants

In addition to identifying just the species of host from the blood meals, we found intraspecific variation in mtDNA haplotypes within host species (Figure 1; Extended data 4^18^). Single haplotypes were found for all of the domestic hosts identified: mouse (*Mus musculus*), chickens (*Gallus gallus*), goat (*Capra hircus*) and cattle (*Bos taurus*). Three human haplotypes were identified, with the majority showing similarity to cytb sequences identified from tsetse blood meals in the Serengeti, Tanzania (type 2; n = 21) or Zambia (type 3; n = 3) (Extended data 4^18^). However, three samples with evidence of only a single host matched an Asian haplotype from Taiwan (type 1: one from each of the three sites), which is the ethnic origin of the primary researcher; these three were excluded from analyses because they were suspected laboratory contaminants. Three additional samples were identified as human but the sequences were not clean enough to resolve the haplotype because they were all identified in flies that appear to have fed multiple times. There was extensive variation in haplotype diversity among the wild hosts but this was not always related to their relative abundance in the samples (Figure 1; Table 2).

### Variables influencing tsetse feeding behaviour

A qualitative summary of variation in feeding behaviours (single vs multiple) of flies in relation to their sex, age, sample site and type of host fed on is provided in Figure 2. Although there was variation in the sex and age distribution of flies across sites (Extended data 5^18^), the most striking pattern distinguishing single and multiple feeding was in relation to differences in the type of hosts fed on.

The two sites from the Shimba Hills (Buffalo Ridge and Zungu Luka) showed a higher proportion of flies that appear to have fed on multiple hosts than the site from Nguruman (Mukinyo): 41.8% from Buffalo Ridge; 61.5% from Zungu Luka, compared with 2.0% from Mukinyo. However, this appeared to be influenced by the type of host (Table 2; Figure 2). Buffalo Ridge showed a predominance of flies that had fed on wild hosts (65.5%) and most individuals with a dominant domestic or human host had fed on multiple species (18/19, compared with 5/36 for wild hosts). Cloning revealed that all five of the individuals classified as multiple feeding had fed on humans and at least one other domestic animal; four of the individuals had also fed on a wild host (Extended data 3^18^). In contrast, for Zungu Luka, humans comprised 43% of dominant hosts identified compared with 33% domestic and only 23% wild animals. Similar to Buffalo Ridge, the majority of flies feeding on non-wild hosts fed on more than one host species (23/30), compared with only 1/9 of the flies for which dominant sequences were identified as wild hosts. All three multiple feeding flies cloned from this site had fed on humans, with one also having fed on both domestic (goat, mouse) and wild (antelope) hosts, one on a single wild host (bushbuck) and another on a domestic host (chicken) (Extended data 3^18^). At Mukinyo, 99% of flies had fed on wild hosts, with only a single fly identified as having recently fed on a domestic host (identified as single feeding on cattle) and no human hosts were detected. Moreover, only 3/149 flies with dominant wild hosts had fed on more than one host species (confirmed by cloning for two of the individuals; Extended data 3^18^).

Using the type of feeding behaviour (single vs multiple hosts) as a binary response variable, the final model selected by maximum likelihood included a highly significant effect of type of host (LRT: *χ^2^=* 52.0, df = 2, *p* = 5.09e-12) and a significant effect of site (LRT: *χ^2^=* 13.2, df = 5, *p* = 0.001). Examining the odds ratios (OR) indicated a substantially lower incidence of multiple feeding on wild compared to domestic hosts (OR = 0.009; CI = 0.001-0.045) but similar incidence in humans and domestic hosts (OR = 0.232: CI = 0.031-1.151). As might be expected based on the difference in distribution of hosts, Mukinyo showed lower levels of multiple feeding (OR = 0.081; CI = 0.014-0.323) than Buffalo Ridge, whereas there was little difference between the two Shimba Hill sites (OR = 0.407; CI = 0.081-1.528). Comparing the residual deviance (104.12 on 239 df) and null deviance (247.49 on 243 df) indicated that there was no evidence for over-dispersion and McFadden’s pseudo-R^2^ was 0.58, indicating a relatively good fit to the data that the final model explained.

### Prevalence of *Trypanosoma spp.* in relation to *G. pallidipes* feeding patterns

Across sites, 44% (n = 107) of the flies for which hosts could be identified to species tested positive for trypanosomes with 54% (29/53) associated with dominant domestic hosts and 41% (79/194) with wild (Table 1). Of flies feeding on multiple hosts, 58% tested positive for trypanosomes, compared to 40% that had fed on single hosts, but this was influenced by the higher rate of infection in Zungu Luka (61.5%), where single feeding was rare, compared to in Buffalo Ridge (36%) and Mukinyo (42%). It was more difficult to interpret patterns by host species because of the large differences in their relative abundance (Extended data 6^18^).

As found in our previous study^17^, generalised linear models using trypanosome presence as a response variable were difficult to interpret. All of the interactions considered except for that between feeding pattern and site significantly explained variation in trypanosome detection (p<0.01). However, testing the fit of the final model based on pseudo-R^2^ (0.08) indicated that only a small amount of the variation in trypanosome presence was explained. The residual deviance (303.22 on 225 df) also suggested over dispersion.

For this reason, multivariate ordination analyses were used to visualise associations between variables. Based on MCA analyses, strong correlations among site of *G. pallidipes* collection, host feeding pattern and type of host were apparent in dimension 1 (Figure 4; Extended data 7^18^). In contrast, trypanosome status was resolved primarily along dimensions 2 and 3, as were sex and age of the flies; a positive association was found between trypanosome positive samples and juvenile male flies, while trypanosome negative flies tended to be found in young female flies.

**Figure 3.**
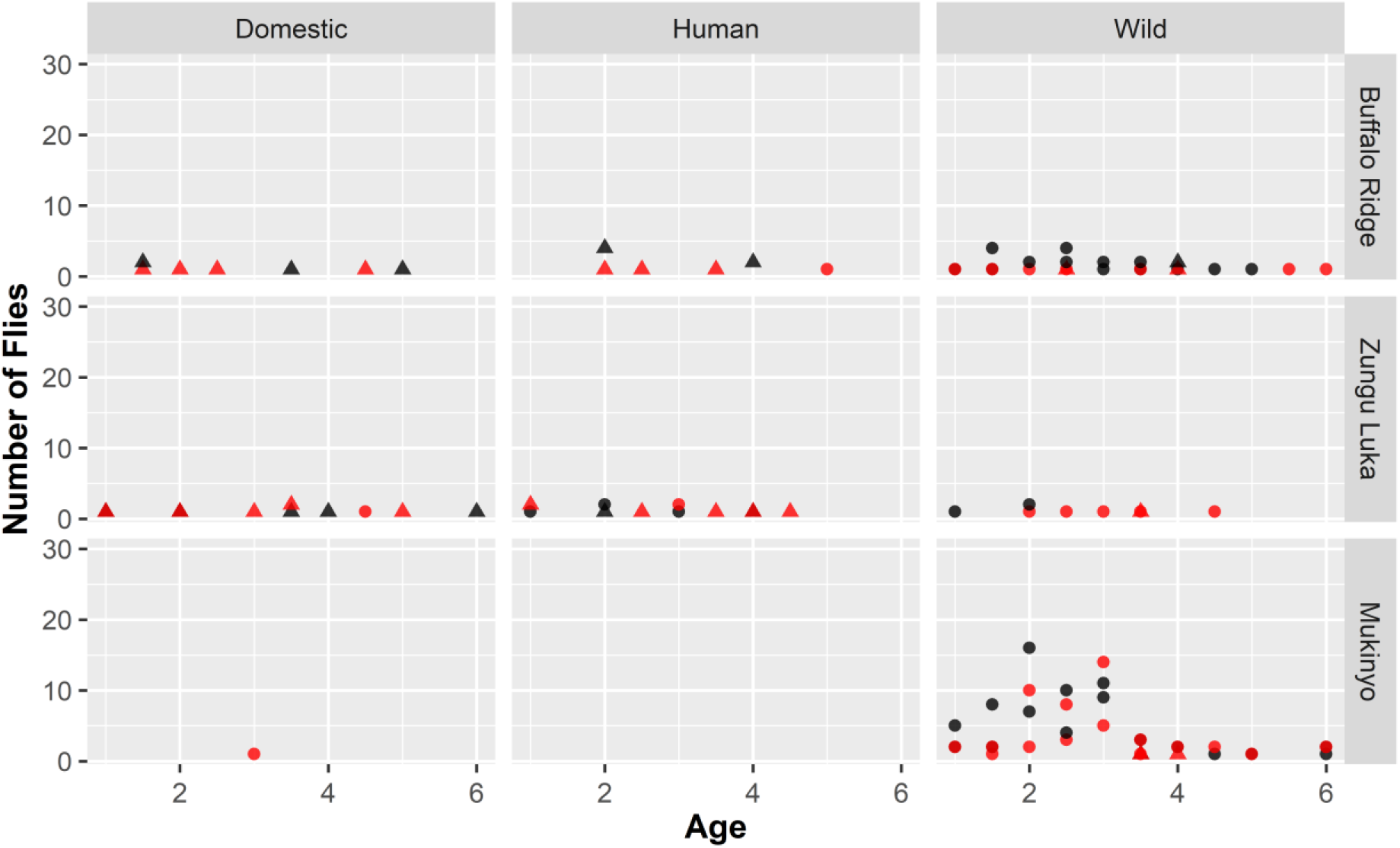
Detection of trypanosomes across sites in relation to fly feeding behaviour, age and site. Trypanosome detection (black = negative, red = positive) is indicated in relation to feeding pattern (circle = single, triangle = multiple), with separate plots by type of host and site. Generalised linear mixed models indicated multiple significant pairwise interactions between feeding behaviours and other tsetse-specific variables. Sex was involved in a significant interaction with type of host but not feeding pattern, but it has been excluded here to more clearly demonstrate the complicated interactions between the other variables.

**Figure 4.**
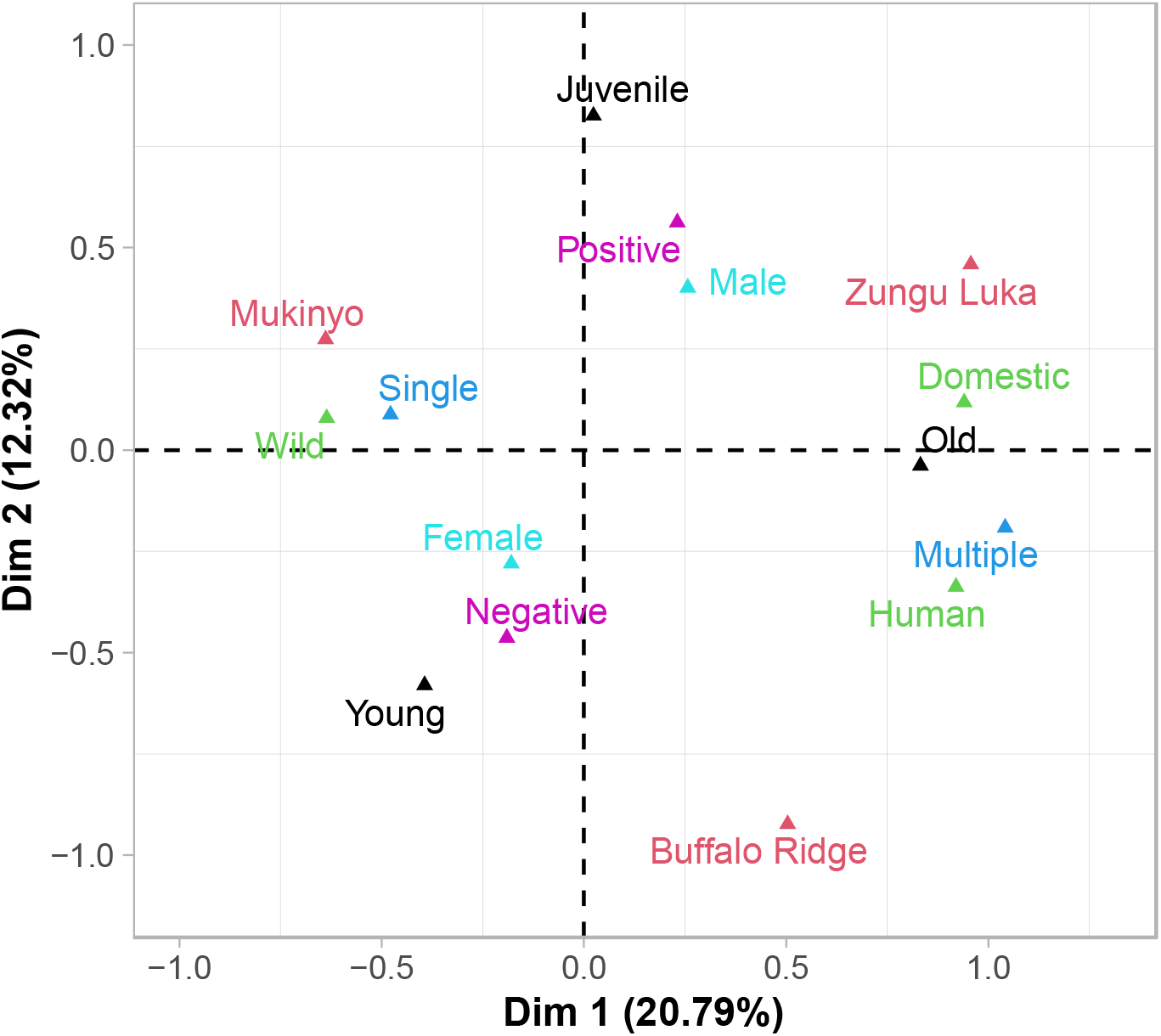
Multiple Correspondence Analysis (MCA), showing associations of dimension 1 (Dim 1; 29.79 % of the variance) and 2 (Dim 2; 12.32% of the variance) in relation to age category (young, juvenile, old), feeding pattern (single or multiple), host type (domestic or wildlife), sex (male or female), site (Buffalo Ridge, Zungu Luka or Mukinyo), and *Trypanosoma spp*. status (positive or negative). This Figure clearly shows the strong association between feeding pattern and host type, driven by the differences in fly behaviour at Mukinyo compared to Zungu Luka resolved along dimension 1. Old flies were also highly correlated with multiple feeding of domestic and human hosts at Zungu Luka Trypanosome status was not explained by variation along dimension 1 but was more related to sex and age of younger flies resolved along dimension 2. Buffalo Ridge was differentiated from the other two populations along both dimensions 1 and 2.

## Discussion

Based on detailed sequence analysis of mitochondrial gene amplicons, our results suggest that individual tsetse flies (*G. pallidipes*) vary markedly in their feeding patterns. In particular, we found that flies feeding on wild hosts tended to show higher feeding success (based on evidence for amplification of only a single host species in blood meals) than those feeding on domestic animals and humans. Although site also influenced patterns of feeding, this was somewhat confounded by the relative abundance of wild hosts that were fed on between the two regions compared. Although previous studies have found a similar diversity of hosts as we found based on analyses using the same cytochrome b primers^40^ or other mtDNA regions^54^, we are not aware of other studies that differentiated single from multiple feeding based on analysis of sequence chromatograms. Moreover, blood meal analyses do not typically assess within-host diversity; our haplotype analysis suggests that there is potential to use feeding arthropods as “flying syringes”^12^ not only for identification of hosts but also could be used to make inferences about host population structure. Our results were not able to clearly test whether host feeding patterns or type of host influenced prevalence of trypanosomes in individual flies. As in our previous study^17^, trypanosome presence was explained by interactions between multiple variables. We had hypothesised that some of this complexity might be reduced by including feeding behaviours, but they also were found to influence variation dependent on other variables. Multivariate analysis using MCA suggested that prevalence of trypanosomes was correlated with sex and age of the flies whereas feeding pattern was correlated with type of host and geographic location.

Together, these results suggest that differences in host communities in different regions could influence the risk of transmission between vectors and hosts in complex ways and highlight the potential for increased transmission risk in interface areas where both livestock and domestic hosts coexist.

### Host diversity in *G. pallidipes* blood meals

We identified the dominant hosts for 46% of the *G. pallidipes* samples screened (56% of the samples that showed positive amplification products), which is comparable or higher than previous studies using the same primers^5,12^. We found extensive variation among the species fed on in two different geographic areas. In the Shimba Hills, where a fenced wildlife protected area is located within a few km of human settlements, flies fed on both domestic and wild hosts, with blood meals from both host types detectable within individual flies. In contrast, in the Nguruman region, only a single fly was identified that had fed on a domestic host. This is consistent with Muturi *et al*.^54^, who also did not identify domestic hosts in their survey of the Nguruman region, despite finding predominantly cattle blood meals at a site surveyed in Uganda. The results from Nguruman may be due to sampling time and the large-scale shifts in cattle grazing sites according to season (Masiga, unpublished). Snow *et al*.^65^ suggested that, even though flies in areas dominated by cattle fed readily on these domestic hosts, a positive correlation between the number of wild herbivores and the abundance of *G. pallidipes* suggested that feeding success was poor on local livestock (based on a low density of flies where cattle were numerous). The use of insecticides on cattle also could help to explain decreased density of tsetse traversing from wildlife-protected areas through to livestock dominated areas in interface areas^48,49^

Surprisingly, no domestic cattle were detected in our study from the Shimba Hills, despite the proximity to settlements with mixed herds of cattle, sheep and goats ^55^. However, domestic hosts were also not identified in the Shimba Hills region in a previous study based on host detection using haemagglutinin assays^65^. This could indicate that flies avoid cattle when more favourable hosts are present. However, there also could be seasonal differences, as trypanosome prevalence in cattle was found to be high (33.9%) in Kwale County, in a previous study that also found *G. pallidipes* at high abundance^51^. At Mukinyo a single individual fed on domestic cattle but there was a very low proportion of flies that fed on multiple hosts (2%) and a predominance of buffalo (71%) among the samples where the dominant host could be identified. It is possible that buffalo are abundant hosts that are easy to feed on and so flies could learn to return to the same host species^14^. Our results in general are consistent with higher feeding success on wild compared to domestic hosts.

Our finding of African buffalo as the main hosts of *G. pallidipes* in Nguruman and Buffalo Ridge supports previous reports that ruminants are attractive to adult *G. pallidipes*, *G. fuscipes* and *G. brevipalpis*^1,30^. However, host selection has been found to vary extensively by population (Extended data 7^18^). Differences across studies could be due to differences in methodology but also could be due to microhabitat differences^66^, such as seasonal variation in host availability, the vegetation type or cover at particular sites, or particular environmental conditions in different years, which affects overlap of habitat and activities between tsetse flies and hosts^1,19^. It is interesting that no buffalo blood meals were detected at Zungu Luka, despite its close proximity (~ 20 km) to Buffalo Ridge, where buffalo are abundant. This could suggest that flies feeding in human settlements move into the park to feed on wildlife but once feeding on their preferred wildlife, they do not move out into the human-settled regions or that flies tend to dwell proximally to where bloodmeals are readily available. It would be interesting to quantify relative abundance of hosts of different types and directionality of movements to test this hypothesis. Specific choice tests between domestic and wild hosts also could reveal important information about preferences that could inform control interventions^70^ as has been done for malaria-carrying mosquitos^50^. Nevertheless, the finding of flies collected in the same traps feeding on both wild and domestic hosts emphasizes the high potential for cross-feeding between these host types when they occur sympatrically.

Humans have been suggested as inappropriate hosts because they camouflage their odours, apply chemical repellents, and react strongly to tsetse bites, which could result in unsuccessful feeding^9,10,29^ that could lead to host switching. Hargrove^29^ found that the presence of humans not only repelled tsetse flies but also inhibited the landing response to approach other potential hosts nearby. Most of the mtDNA haplotypes we identified from human samples were consistent with those expected regionally and one haplotype was shared with previous published sequences from the Serengeti (Auty et al. 2016a; Extended data 4^18^); we also did not find evidence of feeding on humans in Mukinyo. Although measures were taken to rule out contamination, these results are surprising and patterns of tsetse feeding in areas of higher human density should be investigated further.

### Variables influencing tsetse feeding behaviour

We found that the propensity for feeding on single compared to more than one host species was highly influenced by the type of host fed on, with more single feeding on wild hosts than on humans or domestic host. Cloning and sequencing revealed that some flies feeding on domestic or human hosts had fed on up to four different host species and confirmed that single feeding was more common in flies feeding on wildlife. Theoretically, the number of clones could be used to predict which host was last fed on, but this would also depend on the rate of feeding of the fly (e.g. if they were interrupted and switched hosts very rapidly, more than one blood meal might have a similar DNA concentration) and lack of bias in PCR amplifications. There also could be behavioural differences that could result in detection biases: 1) flies might feed more thoroughly on their preferred hosts (such as buffalo), increasing the blood meal volume from that host; 2) flies might feed multiple times on the same host species occurring at high local densities (suggested here by the presence of multiple haplotypes of the same host species in some cases); or 3) hosts might differ in effective defence mechanisms, resulting in low blood meal volumes due to interrupted feeding^68^. If feeding on an initial host is interrupted or too low quality (“unsuccessful”), flies might switch hosts. Unsuccessful feeding of tsetse flies on cattle have been attributed to host defence, such as twitching the skin, flicking the tail, flicking the ears, and kicking or stamping^63^. Wild animals might react less to tsetse flies feeding and/or be surrounded by less other biting insects than domesticated animals. Nevertheless, our results suggest higher host fidelity (or feeding success) when feeding on wild, compared to domestic, hosts.

### Prevalence of Trypanosoma spp

There was not a clear association between prevalence of trypanosomes and type of host or host-feeding patterns in the tsetse flies. In our previous study Channumsin *et al*.^17^, we found that trypanosome prevalence was explained by complicated interactions between age, sex and sampling site of the tsetse flies. Here we found that detection of trypanosomes was also significantly influenced by interactions with these tsetse-specific variables with both host type and feeding patterns. This made it difficult to test our hypothesis that the tsetse feeding behaviour might explain some of the variation in trypanosome detection. Specifically, we hypothesised that feeding on multiple hosts could increase risk of trypanosome infection in flies. However, this was not apparent in the multivariate analysis using MCA (Figure 4) suggested a stronger correlation among feeding pattern, host type and site than with trypanosome status, sex and age of tsetse flies. The blood meals we analysed also only reflect the most recent feeds and so likely do not reflect their overall feeding history. Bouyer *et al*.^14^, suggested that repeated feeding on the same host species was likely to increase risk of trypanosome transmission within species, but to decrease risk between species. There is some evidence that trypanosome infection might influence feeding success and feeding behaviour of the flies, but it is not conclusive^45^. For example, high numbers of *T. congolense*, which attach to the cuticle of the proboscis, could interrupt feeding and result in more frequent probing^38^. Alternatively, nutritional status of the flies could affect their relative susceptibility to trypanosome establishment^41^. Thus, an association between the frequency of feeding and trypanosome infection status should be further studied in laboratory experiments to test whether trypanosome infection causes a feeding pattern change or differences in feeding patterns promote trypanosome infection. Nevertheless, our results did not suggest an increased prevalence of trypanosomes in communities where both domestic and wild hosts were fed on that would suggest increased risks in livestock interface regions.

### Amplicon-based blood meal analyses

Although blood meal analyses provide a powerful tool for investigating feeding behaviours of haemotophagous insects, the potential for biases in any PCR-based approach deserves consideration. For example, we found that a higher proportion of hosts could be resolved from Mukinyo than the other areas, which could be due to the dominance of wild hosts but could also be due to higher fidelity of the primers used on the species of hosts detected. Previous studies comparing the relative reliability of cytb and COI mtDNA e.g. Muturi *et al.*^54^ have found that neither alone amplifies products from all potential host types present. In the Shimba Hills, although goats were identified from flies sampled from both sites, there were also additional samples that matched goats in BLAST analyses that were not included in the analysis because it was difficult to determine whether the sequences represented multiple feeding or just poor sequence quality. Moreover, blood meal analyses rely on completeness of reference databases. We found several cytochrome b haplotypes that were closest to antelope in BLAST but the similarity was too low to resolve to species (93%); this lack of reference sequences could have led to underestimates of host usage in previous blood meal analyses. Analyses of blood meals also do not typically consider the possibility of amplification of nuclear copies of mitochondrial genes (numts), the presence of which can vary dramatically across vertebrate species^32^. It was for these reasons that we took a conservative approach to interpreting feeding patterns based on blood meals by only considering sequences where the dominant host could be clearly identified by direct sequencing (or cloning), While this meant that we likely underestimated the rate of feeding on multiple host species, a clear pattern remained that fewer ambiguous sequences were found at Mukinyo, where wild hosts dominated, than at the other sites (26% vs 44%, respectively).

There has been a recent shift towards using deep sequencing approaches for amplicon-based host identification^35,62^, which would allow more rigorous testing of potential biases and could also allow simultaneous targeting of hosts and trypanosomes by using multiplexed approaches^23^. Non-PCR based assays such as high-resolution melting point analysis have already shown high promise as alternatives for blood meal analysis^22,59^. However, deep sequencing following enrichment approaches rather than PCR amplification, such as hybrid sequence capture^7,53^, have the potential to not only provide a more comprehensive analysis of host diversity, but could allow clearer interpretation of relative read numbers in relation to feeding patterns.

## Conclusions

Identification of the hosts that *G. pallidipes* fed on based on direct PCR sequencing revealed evidence for both use of a wide range of hosts and multiple feeding bouts by individual flies. However, in wildlife dominated areas, there was a much stronger tendency for flies to feed on single host species compared to sites where domestic hosts were more commonly fed on, with individual flies feeding on up to four different detectable host species. If this indicates that domestic animals are not preferred hosts, this could have important implications for understanding risk of transmission of trypanosomes between wildlife and livestock in interface areas. Our results also demonstrate the value of detailed sequence analysis of blood meals of haematophagous insects to include not only identification of the host species but patterns of feeding by individual flies in relation to their sex, age and habitat. The increased accessibility of deep sequencing approaches opens up new possibilities for more detailed assessments, which might also include the ability to predict the timing or success of feeding on different hosts based on relative read depths.

## Data Availability

Sequences have been deposited to Genbank, with accession numbers MN148732-MN148768 (Extended data 4^18^).

## Acknowledgments

This work was funded by the Wellcome Trust (grant number: 093692) to the University of Glasgow. For the purpose of Open Access, the author has applied a CC BY public copyright licence to any Author Accepted Manuscript version arising from this submission. The authors are grateful to the Kenya Wildlife Service for permission to perform this study in protected areas. Esther Waweru and James Kabii provided technical and administrative assistance at *icipe*. James Kabii, Moses Tanju, Loya Suleyman, John Andoke (deceased) and his team (*icipe*) provided invaluable expertise and help for the trapping of tsetse. We thank DNA Sequencing & Services (MRC I PPU, School of Life Sciences, University of Dundee, Scotland, https://www.dnaseq.co.uk) for DNA sequencing. The comments from three anonymous reviewers on a previous draft substantially improved the manuscript.

